# Acvr2b receptors dose-dependently regulate dorsoventral patterning, FOP ACVR1-R206H signaling, and limit BMP/Nodal signaling

**DOI:** 10.1101/2025.10.07.680911

**Authors:** Jeet H. Patel, Benjamin Tajer, Jira White, Finn Warrick, Mary C. Mullins

## Abstract

BMP signaling drives dorsoventral (DV) axial patterning in vertebrates and invertebrates, with BMP dimers recruiting tetrameric receptor complexes to phosphorylate SMADs that activate target gene expression. In zebrafish DV patterning, Bmp2/7 heterodimers exclusively signal, assembling a receptor complex of two distinct type I receptors, Bmpr1 and Acvr1l, that canonically bind Bmp2 and Bmp7 ligands, respectively. Here, we tested if the two distinct classes of BMP type II receptors, Bmpr2 and Acvr2, also act in the gastrula signaling complex. We mutated all four *acvr2a* and *acvr2b* genes in the zebrafish and found maternal-zygotic depletion of Acvr2b receptors causes severe embryonic dorsalization, while *bmpr2* genes are not expressed in the gastrula, indicating Acvr2b is the primary type II receptor transducing the BMP signal. Further, while phosphorylated Smad5 (pSmad5) was reduced throughout the early gastrula, unexpectedly, pSmad5 was nearly abrogated at the embryonic margin, where Nodal signaling is active in specifying the mesendoderm. We found that inhibiting Nodal signaling by depleting its co-receptor, Oep, in MZ-Acvr2b deficient embryos restored marginal pSmad5 to non-marginal levels, suggesting that Nodal competes with BMP for Acvr2a receptors at the margin. Further, we find that while Bmpr2 is not expressed in the gastrula, it can signal in the absence of Acvr2b, indicating that, unlike the type I receptor classes, the type II receptor classes do not uniquely function in the Bmp2/7 signaling complex. We further demonstrate that the ACVR1-R206H Fibrodysplasia Ossificans Progressiva human disease-causing mutant receptor, ACVR1-R206H, displays an increased sensitivity to Acvr2b dosage compared to wild-type Acvr1l, indicating that Acvr2b receptor availability restricts multiple modes of signaling in the gastrula.

## INTRODUCTION

BMP signaling functions as a major pathway in embryonic development, regulating patterning of the dorsoventral (DV) embryonic axis, neural tube, and digits in addition to vascular development and angiogenesis (Chen et al., 2004; Wang et al., 2014; Yan and Wang, 2021; Zinski et al., 2018). The general mechanism of BMP signaling is well defined. BMP ligands are secreted as dimers, binding a serine-threonine kinase receptor tetramer consisting of two type I and two type II receptors. Upon ligand binding, the constitutively active type II receptor phosphorylates the serine/threonine rich GS domain of the type I receptor, activating it. Active type I receptors phosphorylate intracellular SMAD transcription factors, which then accumulate in the nucleus and regulate target gene expression (Chen et al., 2004; Mueller and Nickel, 2012; Yadin et al., 2016).

In zebrafish gastrula, Bmp2/7 heterodimers signal through a heteromeric receptor complex consisting of the type I receptors, Acvr1l and Bmpr1a, and two type II receptors (Bauer et al., 2001; Little and Mullins, 2009; Mintzer et al., 2001; Mullins et al., 1996; Tajer et al., 2021). Interestingly, these two type I receptors have been sub-functionalized in signaling in the gastrula, with Acvr1l kinase activity essential for signaling, while Bmpr1a kinase activity is dispensable (Tajer et al., 2021). Ligand heterodimers and heteromeric type I receptor complexes also function in other BMP signaling contexts (Kim et al., 2019; Nishimatsu and Thomsen, 1998; Peng et al., 2013; Schmid et al., 2000; Suzuki et al., 1997, reviewed in Patel and Mullins, 2025), though it remains unclear why such complexes signal exclusively or outperform homomeric complexes (Karim et al., 2021). To further determine how heterodimers and heteromeric type I receptor complexes signal, it is critical to understand which type II receptors regulate signaling in different contexts.

The role played by the BMP type II receptors, Acvr2 and Bmpr2, in transducing the Bmp2/7 heterodimer signal in zebrafish axial patterning is not known. It has been postulated that two classes of type II receptor may act in DV patterning, parallelling the requirement for the two classes of type I receptor, to confer the subfunctionalized kinase activities of Acvr1l and Bmpr1a (Patel and Mullins, 2025; Tajer et al., 2021). The two distinct classes of type II receptors, Acvr2 and Bmpr2, are represented by six genes in the zebrafish: Acvr2 receptors (*acvr2aa*, *acvr2ab*, *acvr2ba*, *acvr2bb*) that mediate signaling by BMP, Nodal, and TGF-β ligands broadly; and the BMP specific Bmpr2 receptors (*bmpr2a* and *bmpr2b*). Morpholino (MO) or *acvr2* quadruple F0 CRISPR knockdowns do not affect BMP signaling in early gastrulation stages, instead causing later, posterior trunk and tail BMP loss-of-function phenotypes (Albertson et al., 2005; Preiß et al., 2022). This indicates that additional type II receptors transduce BMP signaling during DV patterning. One possibility is that Bmpr2 receptors also transduce the BMP signal; however, MO knockdown and double zygotic *bmpr2a^−/−^*;*bmpr2b^−/−^*mutants do not display axial patterning defects (Monteiro et al., 2008; Zhang et al., 2020). As such, maternally-supplied Bmpr2 and/or Acvr2 receptors could mediate signaling in early embryonic DV patterning, as does maternal Acvr1l, Smad5, and Bmp1a (Kramer et al., 2002; Mintzer et al., 2001; Tuazon et al., 2020).

In the mouse, the three type II receptors, ACVR2A, ACVR2B, and BMPR2, are required in early development. *Bmpr2* mouse mutants are embryonic lethal by E9.5, arresting prior to gastrulation (Beppu et al., 2000), similar to *Bmp4*, *Acvr1*, and *Bmpr1* mutants, indicating a requirement for BMPR2 in early BMP signaling (Gu et al., 1999; Mishina et al., 1995; Winnier et al., 1995). Single mutants of *Acvr2a* or *Acvr2b* do not exhibit embryonic phenotypes (Matzuk et al., 1995; Oh and Li, 1997). However, *Acvr2a^−/−^*;*Acvr2b^−/−^* double mutants arrest development by E8.5, indicating that they function redundantly in early mouse development (Song et al., 1999). As both *Bmpr2* and double *Acvr2a^−/−^*;*Acvr2b^−/−^*mutants display early embryonic lethality, these two classes of type II receptor are not redundant with each other at these stages in mice. As the embryos arrest prior to DV patterning, a specific role for either type II receptor class has not been demonstrated in gastrula DV patterning.

Defining specific requirements for type II receptors in signaling has key implications for modeling of the autosomal dominant disease known as fibrodysplasia ossificans progressiva (FOP). Over 90% of FOP patients have a recurrent mutation in the GS domain of ACVR1, R206H (Hüning and Gillessen-Kaesbach, 2014; Shore et al., 2006), that hyperactivates BMP signaling due to increased BMP and novel Activin responsiveness (Hatsell et al., 2015; Hildebrand et al., 2017). ACVR1-R206H expression in zebrafish embryos causes embryonic ventralization and increased BMP signaling via Smad5 (Allen et al., 2020; Mucha et al., 2018; Shen et al., 2009). Interestingly, FOP mutant receptors display ligand-independent activity (Allen et al., 2020; Hildebrand et al., 2017; Shen et al., 2009), though still require type II receptors for signaling (Bagarova et al., 2013; Lalonde et al., 2023; Le and Wharton, 2012). In the mouse, ACVR2A and BMPR2 act partially redundantly in FOP hyperactive signaling (Bagarova et al., 2013). In zebrafish, MO knockdown of Acvr2ba and Acvr2bb, but not of Acvr2a or Bmpr2, reduced R206H mediated signaling, though did not block it (Lalonde et al., 2023).

Here, we interrogate type II BMP receptor function in DV axial patterning and FOP signaling in the zebrafish. We find that *bmpr2* is not expressed strongly, if at all, prior to gastrulation, leaving only Acvr2a and Acvr2b as possible type II receptors acting in both BMP and Nodal signaling, which specifies the mesendoderm. Examining mutant alleles for *acvr2aa*, *acvr2ab*, *acvr2ba*, and *acvr2bb*, we identify both maternal and zygotic contributions of each gene to embryonic patterning. Zygotic loss of the *acvr2* genes leads to minor dorsalization. Strikingly, maternal and zygotic depletion of only *acvr2ba* and *acvr2bb* (MZ-Acvr2b depletion) dramatically reduces BMP signaling, with near complete loss of phosphorylated Smad5 (pSmad5) resulting in fully dorsalized embryos. Interestingly, we find that inhibition of Nodal signaling by depleting its co-receptor Oep in MZ-Acvr2b deficient embryos partially rescues margin specific pSmad5 loss, suggesting these pathways compete for limited Acvr2a receptors. Further we find that Bmpr2 can functionally substitute for Acvr2b in DV patterning. Finally, we show that signaling by ACVR1-R206H requires a higher Acvr2b dose than endogenous BMP signaling in DV axial patterning.

## RESULTS

### Bmpr2 is not expressed in the early embryo

To determine if different classes of type II receptor signal in the embryo, we first investigated which type II BMP receptors were expressed during DV patterning. All four *acvr2* genes are expressed at the 2-cell stage (maternal deposition) and through shield stage, based on both *in situ* hybridization (Preiß et al., 2022) and RNA-seq data (White et al., 2017) (Figure 1A). Previous *in situ* hybridization analysis found that *bmpr2a* and *bmpr2b* were expressed in embryos ranging from the one-cell stage through 52 hours post fertilization (Monteiro et al., 2008). However, neither *bmpr2* transcript was detected in early stage transcriptomic data sets (White et al., 2017) (Figure 1A). To address this discrepancy, we performed *in situ* hybridization for *bmpr2a* and *bmpr2b* at both early gastrula and 10-somite stages (Figure 1B-E). At 10-somites, both *bmpr2a* and *bmpr2b* are expressed in the head and tail, with *bmpr2b* also detected in the proctodeum (Figure 1C,E), in agreement with prior reports (Monteiro et al., 2008). However, we did not detect *bmpr2a* or *bmpr2b* at early gastrulation, in agreement with the RNA-seq data (Figure 1A-C). We conclude that Bmpr2 is not expressed in the early embryo and therefore cannot participate in axial patterning.

**Figure 1:**
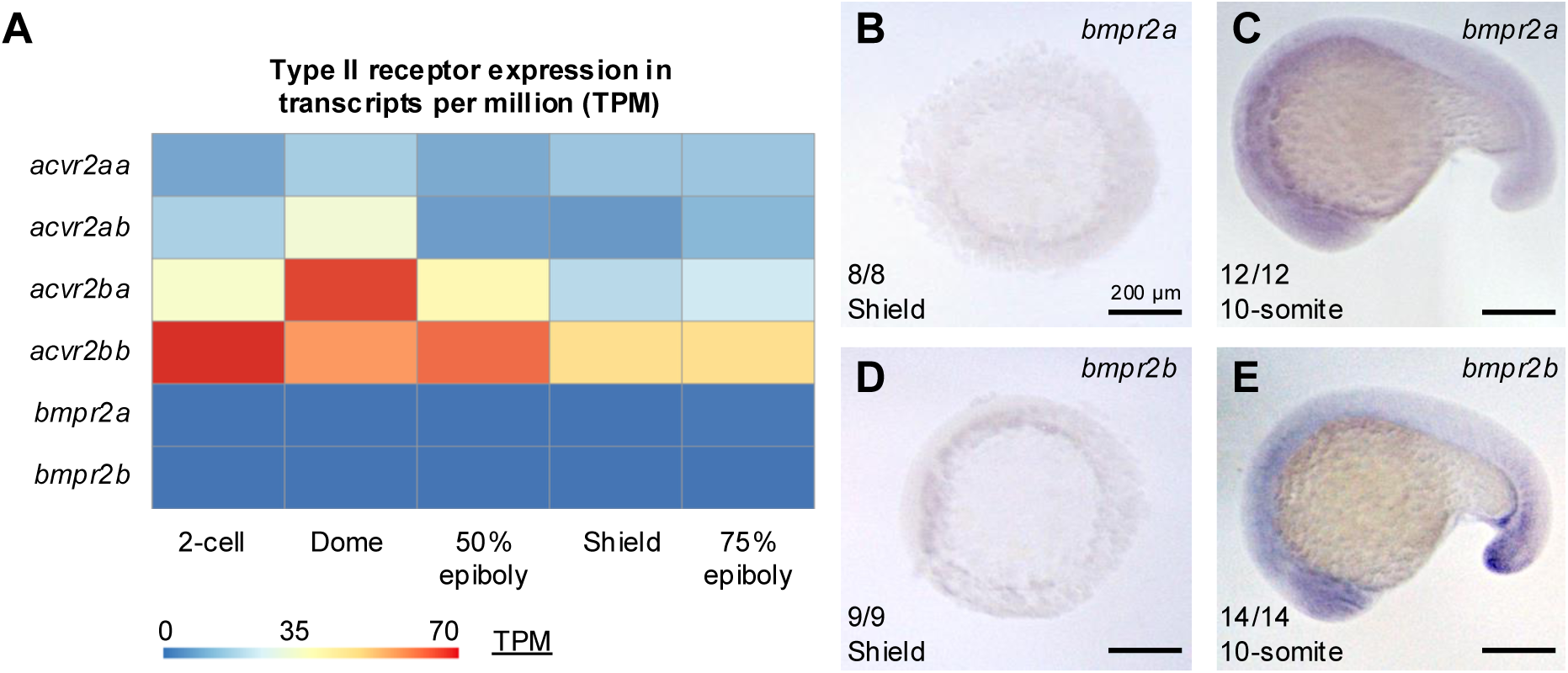
Bmpr2 receptors are not expressed in the early embryo. A) Expression of type II receptor genes in transcripts per million (TPM) for all six type II receptors genes ranging from 2-cell (exclusively maternal) to 75% epiboly, when axial patterning is underway (White et. al, 2017). B-E) *in situ* hybridization at shield (B,D) and 10-somite (C,E) for *bmpr2a* (B,C) *bmpr2b* (D,E).Number of embryos displaying the observed phenotype of the total assayed, representative of 2 experiments.

### Maternal loss of Acvr2b enhances zygotic loss-of-function phenotypes

Since *bmpr2* is not expressed during stages of DV axial patterning, we next asked if maternal *acvr2* expression could account for the normal BMP signaling observed in gastrula stage zygotic *acvr2* CRISPants/mutants (Preiß et al., 2022). We generated mutant alleles of *acvr2aa*, *acvr2ba*, and *acvr2bb* and combined them with the *acvr2ab^sa18285^* allele. All 4 alleles cause predicted frameshifts, terminating in premature stop codons (Supplemental Figure S1). We generated double, triple, and quadruple mutant alleles to test the maternal and zygotic contributions of each gene to embryonic patterning. Phenotypes were assessed at 1 day post fertilization (dpf) based on established dorsalized staging criteria, blind to genotypes (Figure 2A) (Mullins et al., 1996). We then genotyped all phenotypically mutant embryos individually and, as most embryos from these crosses were phenotypically wild type, we did not always genotype each wild-type embryo.

**Figure 2:**
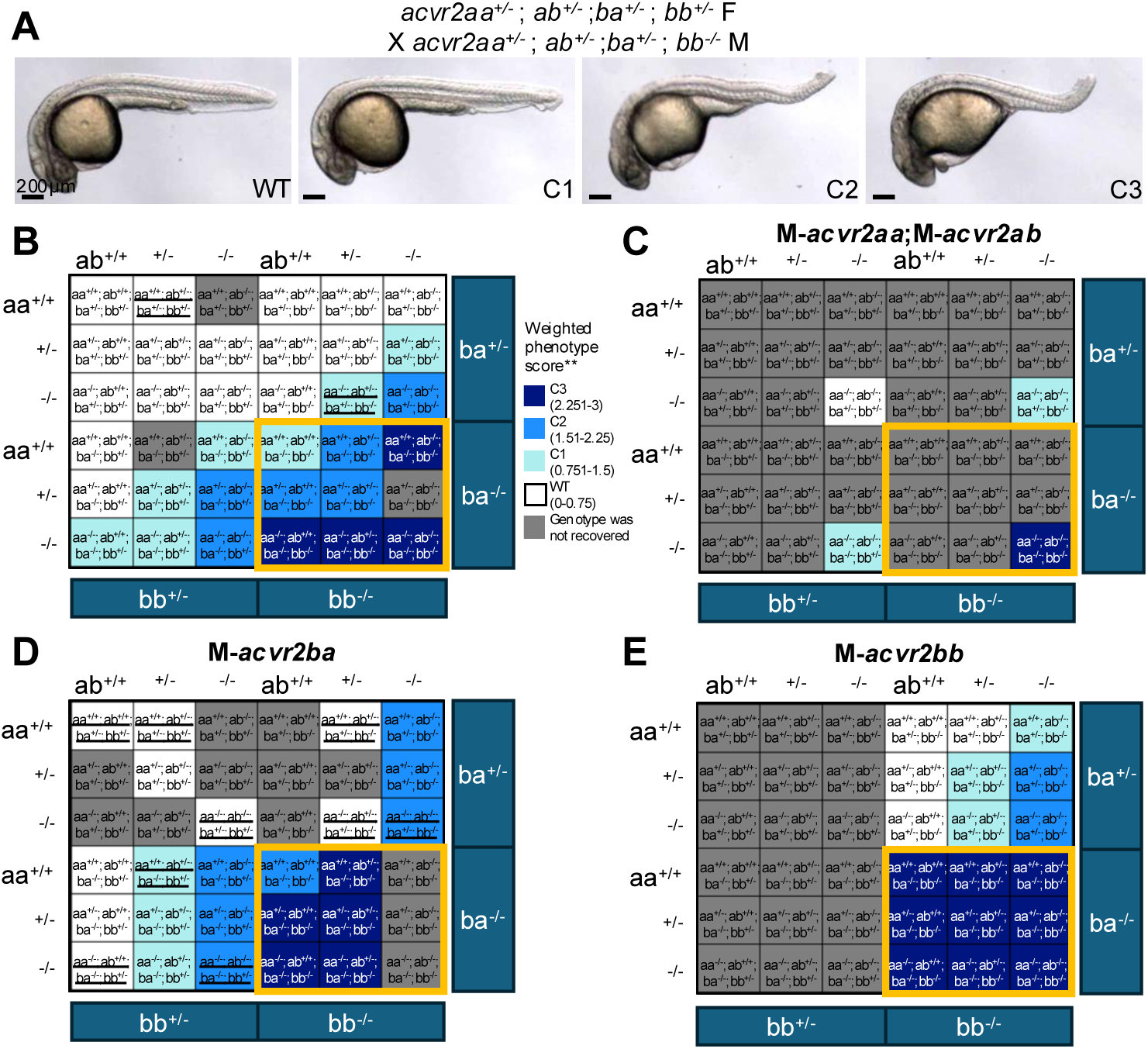
Maternal loss of Acvr2b enhances zygotic embryonic dorsalization. A) Phenotypes observed in Acvr2 mutants based on established dorsalization criteria at 1 dpf. Compared to WT, C1 have a reduced ventral tail fin, C2 have complete loss of ventral tail fin and ventral tail vein, and C3 display C2 phenotypes and fail to extend their yolk extension. B-E) Weighted dorsalization scores for each genotype based on assayed embryos. **Not every phenotypically wild-type embryo was assayed, so in some cases these scores are skewed by low n, full data in Supplemental Figure 2. Phenotypically WT embryos are scored 0, with dorsalized C1, C2, and C3 embryos scored as 1, 2, or 3, respectively. Colors are binned by average phenotype score and grey indicates no embryos of such genotype. Maternal genotypes were: no homozygous maternal mutations (B), M-*acvr2aa^−/−^*;M-*acvr2ab^−/−^* (C), M-*acvr2ba^−/−^* (D) M-*acvr2bb^−/−^* (E). Orange boxes highlight *acvr2ba^−/−^;acvr2bb^−/−^* double mutants. Underlined genotypes indicate only 1 individual was recovered for a genotype. Parental genotypes for crosses are as follows (female X male genotype): B) *acvr2aa^+/−^*;*ab^+/+^;ba^+/−^*;*bb^+/−^* X *acvr2aa^+/−^*;*ab^+/+^;ba^+/−^*;*bb^+/−^*; *acvr2aa^+/−^*;*ab^+/−^;ba^+/−^*;*bb^+/−^* X *acvr2aa^+/−^*;*ab^+/−^;ba^+/−^*;*bb^+/−^*; *acvr2aa^+/−^*;*ab^+/−^;ba^+/−^*;*bb^+/−^* X *acvr2aa^+/−^*;*ab^+/−^;ba^+/−^*;*bb^−/−^* OR *acvr2aa^+/−^*;*ab^+/−^;ba^+/−^*;*bb^+/−^* X *acvr2aa^−/−^*;*ab^−/−^;ba^+/−^*;*bb^+/−^*; C) *acvr2aa^−/−^*;*ab^−/−^;ba^+/−^*;*bb^+/−^* X *acvr2aa^−/−^*;*ab^−/−^;ba^+/−^*;*bb^+/−^*; D) *acvr2aa^+/−^*;*ab^+/−^;ba^−/−^*;*bb^+/−^* X *acvr2aa^+/−^*;*ab^+/−^;ba^−/−^*;*bb^+/−^* OR *acvr2aa^+/−^*;*ab^−/−^;ba^+/−^*;*bb^−/−^*; *acvr2aa^+/−^*;*ab^+/+^;ba^−/−^*;*bb^+/−^* X *acvr2aa^+/−^*;*ab^+/−^;ba^+/−^*;*bb^−/−^*; *acvr2aa^+/−^*;*ab^+/−^;ba^−/−^*;*bb^+/+^* X *acvr2aa^+/−^*;*ab^+/−^;ba^−/−^*;*bb^+/+^*; E) *acvr2aa^+/−^*;*ab^+/−^;ba^+/−^*;*bb^−/−^* X *acvr2aa^+/−^*;*ab^+/−^;ba^+/−^*;*bb^−/−^*;

We found that multiple *acvr2* receptors must be mutant to cause embryonic dorsalization (Figure 2B, Supplemental Figure S2). Single and double zygotic mutants for *acvr2aa*, *acvr2ab*, and *acvr2ba* were mostly wild type, with weak dorsalization when coupled with *acvr2bb*^+/−^ (Figure 2B, middle 3 columns). Triple *acvr2aa^−/−^*;*acvr2ab^−/−^*;a*cvr2ba^−/−^*zygotic mutants displayed weak dorsalization (C1-C2), with moderate C3 phenotypes seen in quadruple mutants (Figure 2B, lower right box). *acvr2bb^−/−^* single mutants were wild type but resulted in minor dorsalization with increasing numbers of other mutated *acvr2* genes (Figure 2B, right 3 columns), indicating that Acvr2 receptors are redundant for BMP signaling and regulate patterning in a gene dosage dependent manner. Notably, the strongest phenotypes were generally observed in zygotic *acvr2ba^−/−^*;a*cvr2bb^−/−^*embryos, consistent with Acvr2b receptors contributing more strongly to signaling (Figure 2B, orange border).

We next investigated if maternal loss of Acvr2 receptors enhanced zygotic mutant patterning defects. Surprisingly, loss of both maternal *acvr2aa* <u>and</u> *acvr2ab* (M-*acvr2aa^−/−^*;M-*acvr2ab^−/−^*) did not enhance the zygotic mutant phenotypes (Figure 2B,C). In contrast, M-*acvr2ba^−/−^*<u>or</u> M-*acvr2bb^−/−^* caused stronger dorsalization for most zygotic genotypes (compare Figure 2B to 2D,E). Specifically, *acvr2ba^−/−^*;*acvr2bb^−/−^*double zygotic mutant embryos with maternal loss of either *acvr2b* gene displayed stronger, C3 dorsalized phenotypes regardless of other mutations (Figure 2D,E, orange border). These findings indicate that maternal loss of Acvr2b receptors increases the severity of phenotypes, supporting a dosage dependent signaling model and suggesting a significant contribution of maternal Acvr2ba and Acvr2bb to early BMP signaling in the embryo.

*Acvr2ba and Acvr2bb predominantly transduce BMP signaling in DV patterning* Maternal loss of a single *acvr2b* gene combined with zygotic loss of other *acvr2* genes showed stronger dorsalization, but did not fully dorsalize embryos, indicating other type II receptor sources for signaling. Therefore, we asked if combined maternal and zygotic loss of both Acvr2b receptors could completely abrogate BMP signaling. Zygotic *acvr2ba^−/−^*;*acvr2bb^−/−^*embryos are dorsalized and not viable to adulthood, preventing generation of double maternal-zygotic (MZ-) *acvr2ba^−/−^*;MZ-*acvr2bb^−/−^*embryos. Instead, we crossed maternal mutants of either gene and injected morpholinos (MO) targeting the reciprocal gene to block translation of maternal and zygotic mRNA. MZ-*acvr2bb^−/−^* embryos injected with *acvr2ba* MO exhibited the strongest C5 dorsalization phenotype, indicative of complete loss of BMP signaling (Figure 3A,B,D,F). For the reciprocal experiment, we injected MZ-*acvr2ba^−/−^* embryos with *acvr2bb* MOs (Dogra et al., 2017, see Methods). Knockdown of Acvr2bb in MZ-*acvr2ba^−/−^* embryos also caused complete dorsalization (Figure 3G,H,J,L). To assess the specificity of MO knockdown, we injected mRNA for the maternally mutant gene, finding rescue of C5 dorsalized embryos to WT or a mild C1/C2 phenotype (Figure 3C,E,I,K). These results indicate that both maternal and zygotic contributions of the Acvr2b receptors facilitate DV embryonic axial patterning.

**Figure 3:**
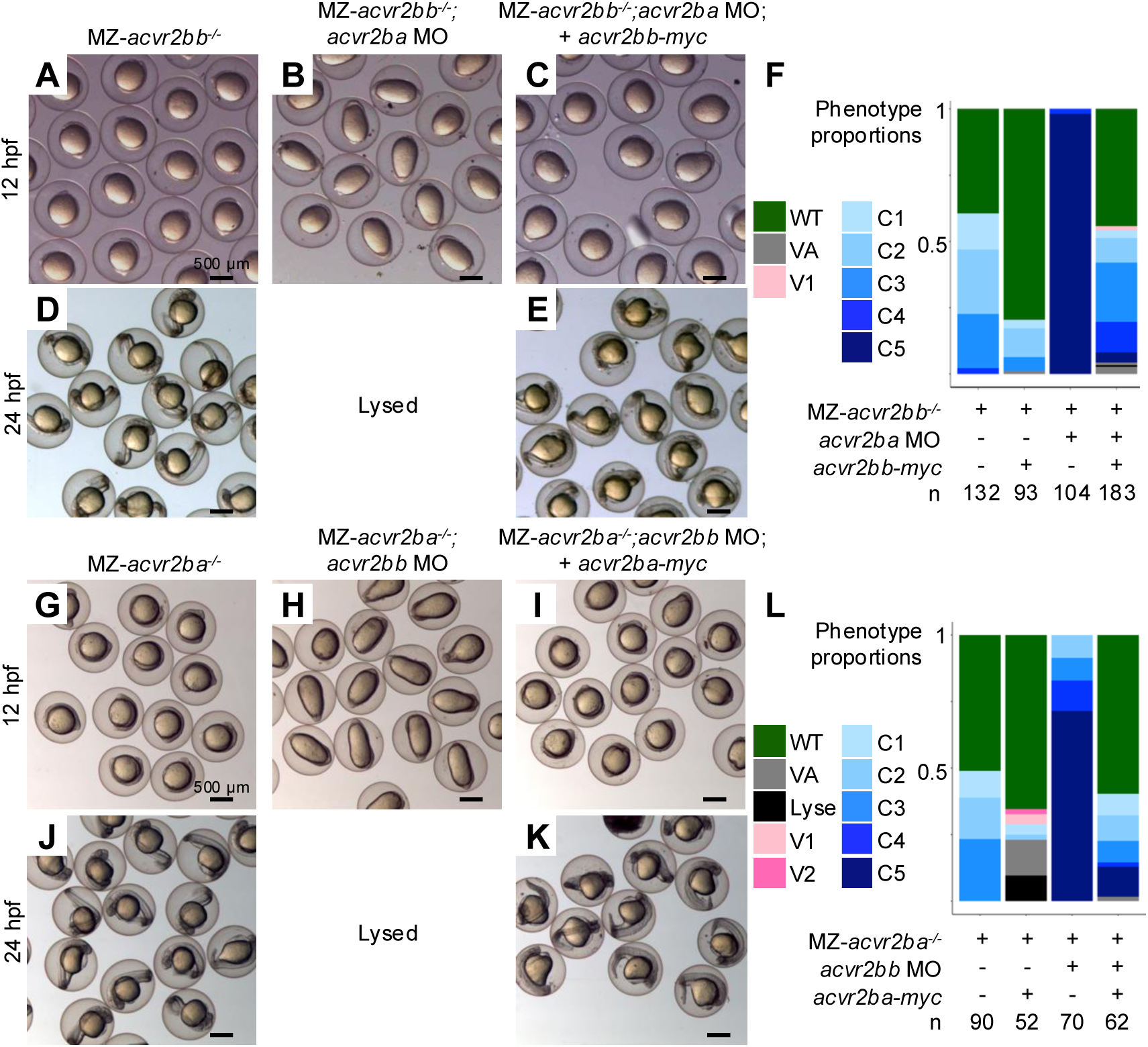
MZ-Acvr2ba and Acvr2bb loss causes complete dorsalization. Representative images of maternal-zygotic (MZ) *acvr2bb* and *acvr2ba* mutants at 12 and 24 hpf (A,D,G,J). Following injection with morpholino for the reciprocal gene (2.5-5ng *acvr2ba* MO or *acvr2bb* MO mix (2.5ng *acvr2bb* UTR MO with 5ng *acvr2bb* ATG MO), embryos are elongated by 12 hpf and lyse, indicative of strong C5 dorsalized phenotypes (B,H). MZ-*acvr2bb* can be rescued with 10-30pg of *acvr2bb* mRNA (C,E) and MZ-*acvr2ba* can be rescued with 50pg *acvr2ba* mRNA (I,K). Head necrosis can be seen in MZ-*acvr2ba* embryos injected with *acvr2bb* MO and rescued (K), likely due to toxicity of the *acvr2bb* MO. Proportions of embryos with different phenotypes in each condition (F,L). Data in F is from 6 independent crosses (except for *acvr2bb-myc* injection into MZ-*acvr2bb* mutants, which is from 3 independent crosses). Data in L is from 3 independent crosses.

### Maternal and zygotic loss of Acvr2ba and Acvr2bb radially expands dorsal cell fates at the expense of ventral fates

To determine if maternal-zygotic loss of Acvr2ba and Acvr2bb abrogates BMP signaling in the gastrula, we examined the expression of DV cell fate markers using *in situ* hybridization. We focused on MZ-*acvr2bb* mutants with *acvr2ba* MO (hereafter MZ-Acvr2b deficient). Depletion of both Acvr2b receptors resulted in circumferential expansion of the dorsal marker *chordin* (*chrd*) (Miller-Bertoglio et al., 1997) and absence of the ventral BMP target gene *sizzled* (*szl*) (Yabe et al., 2003) (Figure 4A,B,D,E), consistent with a complete loss of BMP signaling. *acvr2bb-myc* mRNA rescued *szl* expression and suppressed *chrd* expansion (Figure 4C,F), confirming that dorsalization is caused by Acvr2b loss. To further assess the severity of BMP signal reduction, we examined *bambia* and *foxi1* expression, two direct BMP target genes with lower BMP signaling thresholds than *szl*, using fluorescent *in situ* hybridization (Greenfeld et al., 2021). In shield stage WT embryos, *bambia* and *foxi1* have strong, graded expression across the ventral to dorsal axis, while MZ-Acvr2b deficient embryos have weak, if any, signal (Figure 4G-J). Interestingly, *bambia*, which we previously identified as having the lowest BMP signaling threshold for expression, was expressed weakly in a ventrally restricted domain, suggesting some residual signaling activity in these embryos (Figure 4H). We additionally examined the expression of *pax2.1*, a midbrain-hindbrain boundary marker (Krauss et al., 1992), and *krox20*, a marker of rhombomere 3 at the 1-3 somite stage (Oxtoby and Jowett, 1993). MZ-Acvr2b deficient embryos displayed a radial expansion of these markers, indicative of complete dorsalization of the embryo due to expanded dorsal neurectoderm, which was rescued by wild-type *acvr2bb* expression (Figure 4K-P).

**Figure 4:**
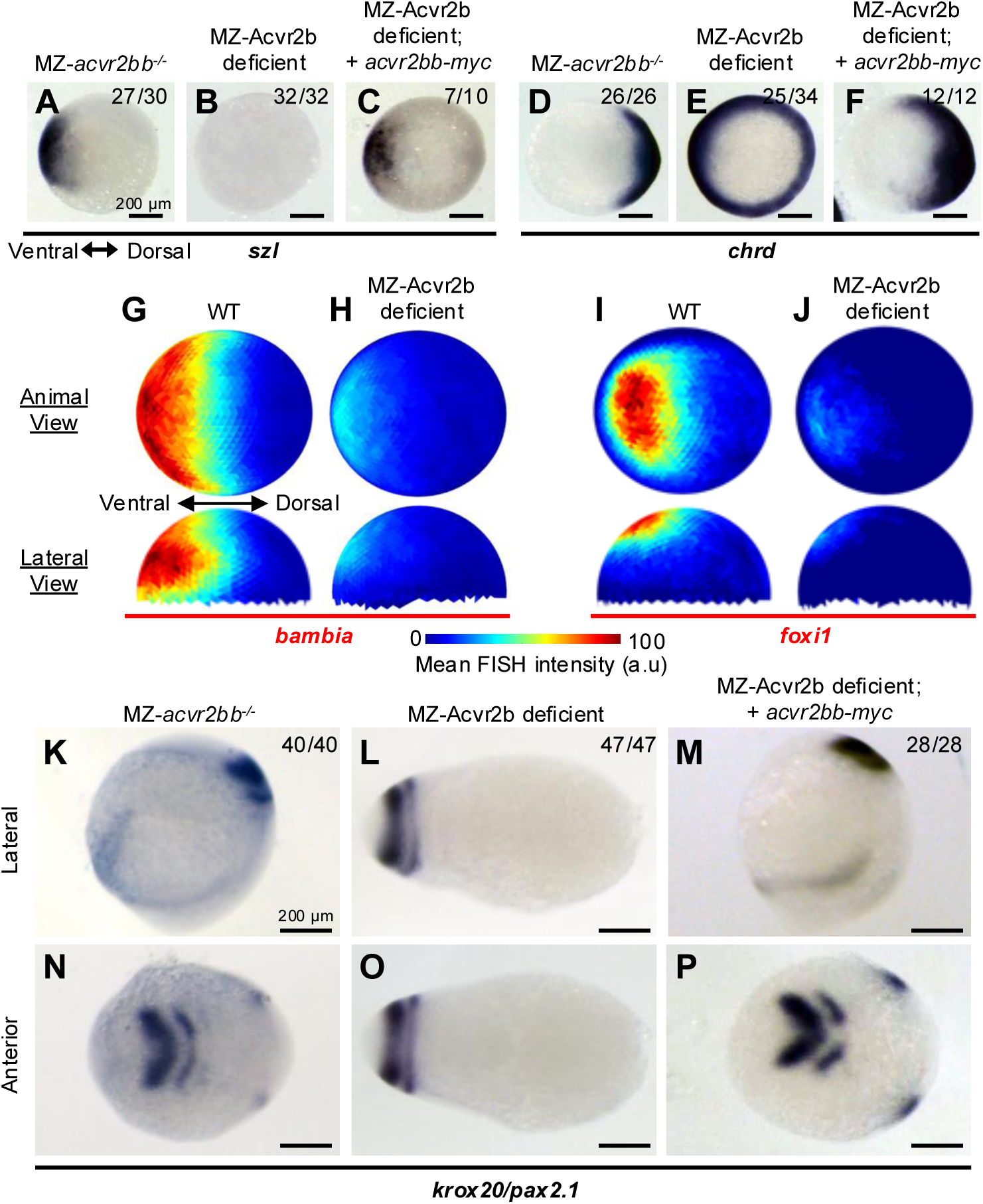
MZ depletion of Acvr2ba/bb causes radial dorsalization of embryos. Representative *in situ* hybridization for *szl* (A-C) *chrd* (D-F) at 60% epiboly stages and *krox20*/*pax2.1* at 1-3 somite stages. *szl* marks the ventral mesoderm (A) and is absent in MZ-Acvr2b deficient embryos (B). *chrd* is expressed dorsally (D) and circumferentially expanded in MZ-Acvr2b deficient embryos (E). 8/34 embryos displayed expanded, but not circumferential *chrd* and 1/34 was phenotypically WT. Both domains are rescued by *acvr2bb* mRNA injection (C, F). Quantification of *bambia* (G,H) and *foxi1* (I,J) TSA *in situ* hybridization at shield stage from. Animal and lateral projections of average signal intensity from WT (G,I) and MZ-Acvr2b deficient (H,J) embryos from 2 independent replicates (n=7 for G,H, n=6 for I, n=5 for J). Lateral view with anterior to top (K,M) and anterior view with posterior to right (N,P) of *pax2.1* marking the mid-hindbrain boundary (anterior stripe) and *krox20* marking rhombomere 3 at 1-3 somite stage (posterior stripe). In L and O, the embryo was rotated 90 degrees as there is no clear lateral, anterior is to left. Circumferential domains for these markers indicates radial expansion of the dorsal domain in MZ-Acvr2b deficient embryos (H,K) that can be rescued with *acvr2bb* mRNA injection (I,L). (K-P) All embryos assayed displayed the phenotype, representative of 3 experiments (2 for C, F, M, and P). Scale bars are 200 μm.

### MZ-Acvr2b deficiency depresses the phosphorylated-Smad5 gradient

To complement characterization of DV patterning changes, we investigated how MZ-Acvr2b depletion alters the gradient of phosphorylated Smad5 (pSmad5). We examined wild type and MZ-Acvr2b deficient embryos at shield stage, performing quantitative analysis of nuclear pSmad5 levels in each cell of the embryo, as previously described (Zinski et al., 2017). Wild-type embryos displayed strong pSmad5 gradients; however, MZ-Acvr2b deficient embryos displayed a significantly reduced pSmad5 gradient across the embryo (Figure 5A-F). In contrast to other BMP signaling mutants, in which the pSmad5 gradient is depressed across the whole embryo (Greenfeld et al., 2021; Tajer et al., 2021; Tuazon et al., 2020), MZ-Acvr2b deficient embryos retain some pSmad5 in more animal regions (Figure 5C,F). While there is a significant, near complete loss of pSmad5 at the embryonic margin, a weaker reduction in the gradient is evident more animally (Figure 5G-H). The weaker pSmad5 animal reduction is consistent with weak *bambia* expression animally (Figure 4H), indicating that BMP signaling is weak, not absent, in animal regions of MZ-Acvr2b deficient embryos. These results taken together demonstrate that maternal and zygotic contributions of Acvr2b receptors (Acvr2ba and Acvr2bb) mediate the majority of BMP signaling required for DV axial patterning, while the remaining Acvr2a receptors are sufficient to partially signal in ventral-animal regions.

**Figure 5:**
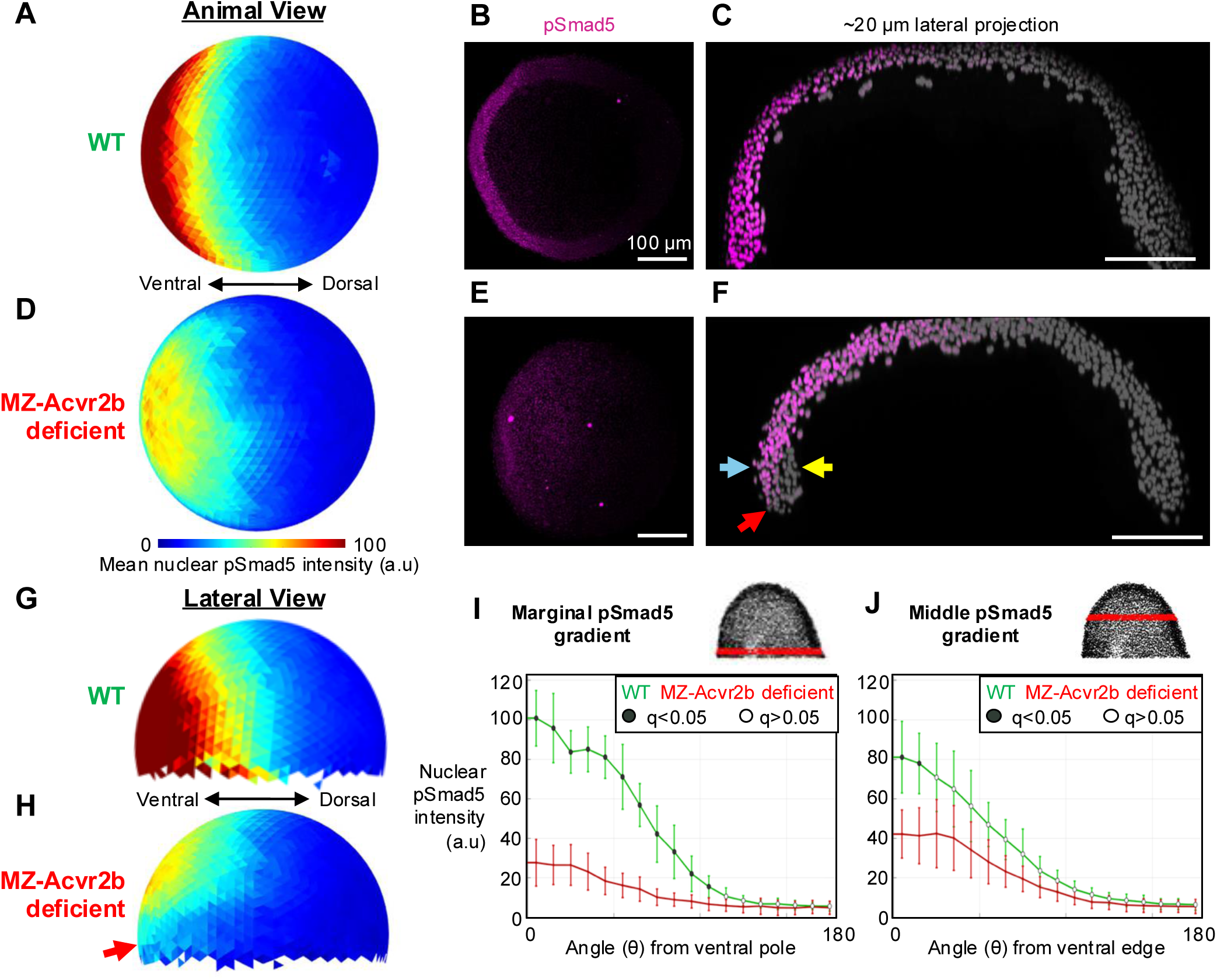
MZ-Acvr2b depletion depresses the pSmad5 gradient most severely at the embryonic margin. Quantification of phosphorylated-Smad5 (pSmad5) in each nucleus from immunostained embryos at shield stage. Animal view projections of average nuclear pSmad5 intensity from WT (A) and MZ-Acvr2b deficient (D) embryos, with 3D-projections of representative embryos (B, E). Lateral projection of Z-stacks to project nuclei through the middle of embryo (C, F) shows differences in animal-vegetal pSmad5 gradients between WT and Acvr2b-deficient embryos. Scale bars are 100 μm. Lateral projections of average nuclear pSmad5 intensity in WT (G) and MZ-Acvr2b deficient (H) embryos. (I, J) Average pSmad5 intensity of a 20 μm band of cells binned at 5° intervals around the embryo at the position marked by the red band above each graph. Black circles indicate significant differences between WT and Acvr2b-deficient embryos in each bin, determined using an unpaired two-tailed t-test on embryo-averaged values, with p-values corrected for multiple testing using the Benjamini–Hochberg procedure (q<0.05). Data are from 2 independent experiments (N=2, n=6 embryos per condition). Red arrows in H and F point to marginal reduction of pSmad5. Yellow arrow in F denotes involuting mesendodermal cells with no pSmad5 signal, while blue arrow points to proximal ectodermal cells.

### Nodal signals at the expense of BMP signaling when Acvr2 receptors are limiting

Strikingly, we observed a complete loss of nuclear pSmad5 in the involuting mesendoderm compared to the ectoderm (Figure 5F, compare yellow arrow to blue arrow). As Acvr2 receptors function in both BMP and Nodal signaling, we hypothesized that the pronounced reduction in marginal BMP signaling could be due to preferred Acvr2 recruitment for Nodal signaling in mesendoderm specification. To test if BMP signaling is limited due to type II receptor signaling through Nodal at the margin, we depleted Nodal signaling and asked if it could restore BMP signaling to levels present more animally in MZ-Acvr2b deficient embryos. We knocked down an obligate co-receptor for Nodal signaling, One-eyed pinhead (Oep) (Gritsman et al., 1999), via morpholino (MO) (Feldman and Stemple, 2001; Nasevicius and Ekker, 2000), and performed quantitative pSmad5 analysis. Interestingly, we found that inhibiting Nodal signaling in MZ-Acvr2b deficient embryos rescued pSmad5 at the ventral embryonic margin (Figure 6) to levels found in more animal cells (Figure 6D-G,I). Since Nodal signaling normally acts only in marginal cells, the pSmad5 gradient in animal cells was not impacted by Oep depletion, as expected (Figure 6D-F,H,J). Thus, marginal cells are uniquely sensitive to Acvr2 receptor availability driving BMP/Nodal signaling.

**Figure 6:**
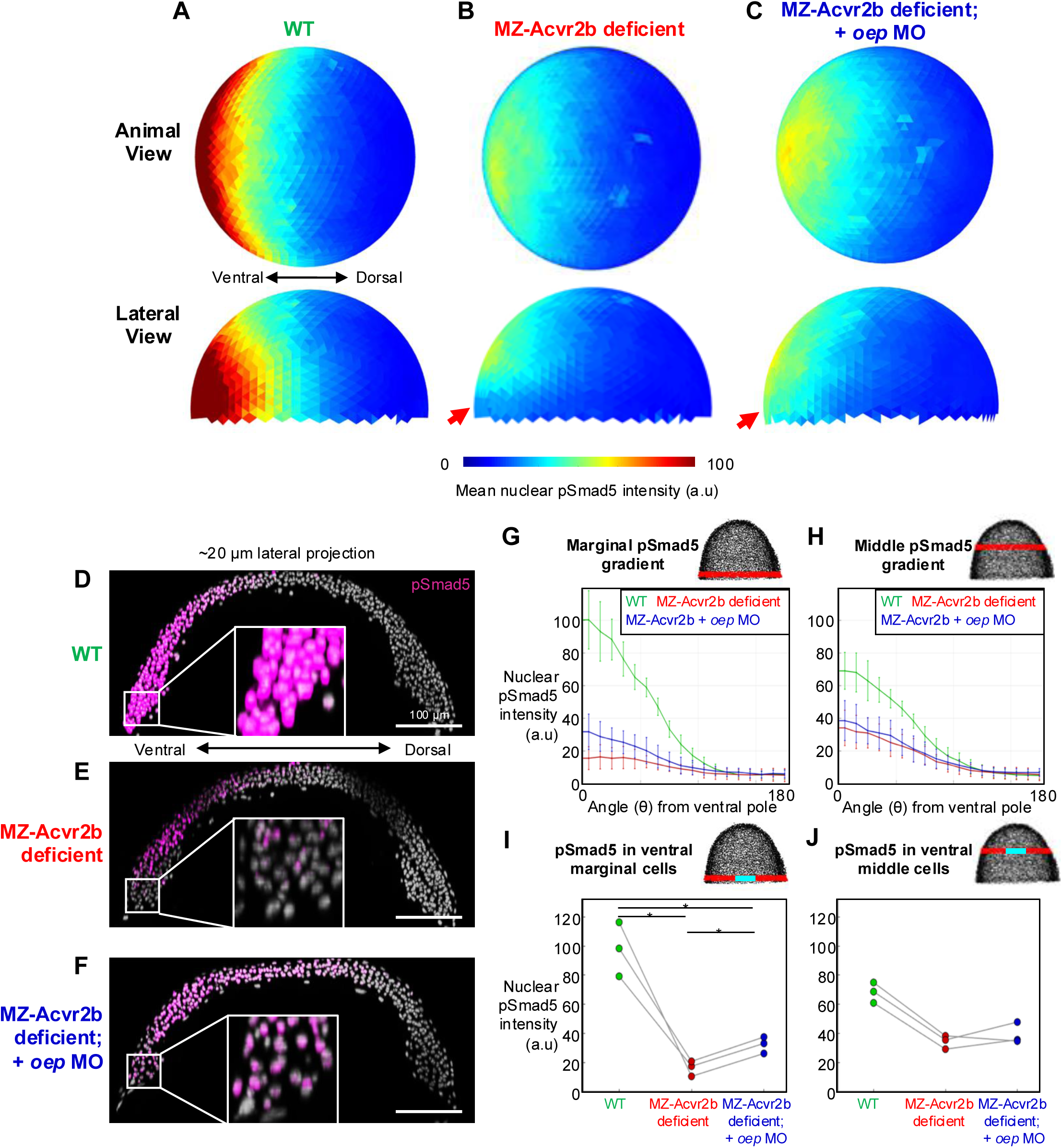
Nodal inhibition restores marginal pSmad5 activity to non-marginal levels. Quantification of phosphorylated-Smad5 (pSmad5) in each nucleus from immunostained embryos at shield stage. Animal and lateral projections of average nuclear pSmad5 intensity from WT (N=3, n=8) (A), MZ-Acvr2b deficient (N=3. n=8) (B), and MZ-Acvr2b deficient with 5ng *oep* MO (N=3, n=11) (C) embryos. Lateral projection of the middle of embryo (D-F). White box marks 50×50 μm zoomed inset in center. Scale bars are 100 μm. (G-H) Average pSmad5 intensity of a 20 μm band of cells binned at 20° intervals around the margin at the position marked by the red band above each graph. (I, J) The ventral most cells (20° region, marked by cyan in schematics) were averaged per embryo. Each point in I-J represents the mean intensity per replicate for each condition, with lines connecting paired replicates. Statistical significance was evaluated using paired two-tailed t-tests with p-values adjusted using the Holm-Bonferroni method. * indicates q<0.05. Red arrows in B and C point to marginal reduction/rescue in pSmad5.

### Bmpr2 can function in place of Acvr2b in DV axial patterning

As the type I receptors Acvr1l and Bmpr1a are not redundant in gastrula BMP signaling (Little and Mullins, 2009; Tajer et al., 2021), we next asked if Acvr2b receptors function uniquely in the heteromeric BMP receptor complex, or if the other type II receptor class, Bmpr2, could transduce BMP signaling in the gastrula. Thus, we tested if Bmpr2b could rescue MZ-Acvr2b deficient embryos. We injected MZ-Acvr2b deficient embryos with mRNA encoding *bmpr2b*, a BMP specific type II receptor, and tested if gastrula DV patterning defects were rescued. We examined early gastrula (60% epiboly) embryos to directly assess if *bmpr2b* was sufficient to restore proper DV patterning. We found that *bmpr2b* expression was sufficient to suppress ventrally-expanded *chrd* expression and restore ventral *szl* expression in MZ-Acvr2b deficient embryos (Figure 7). This indicates that while Bmpr2b is not normally present in the embryo, it can transduce the Bmp2/7 signal in the absence of Acvr2b and that, unlike type I receptors, type II receptors can function redundantly in the gastrula.

**Figure 7:**
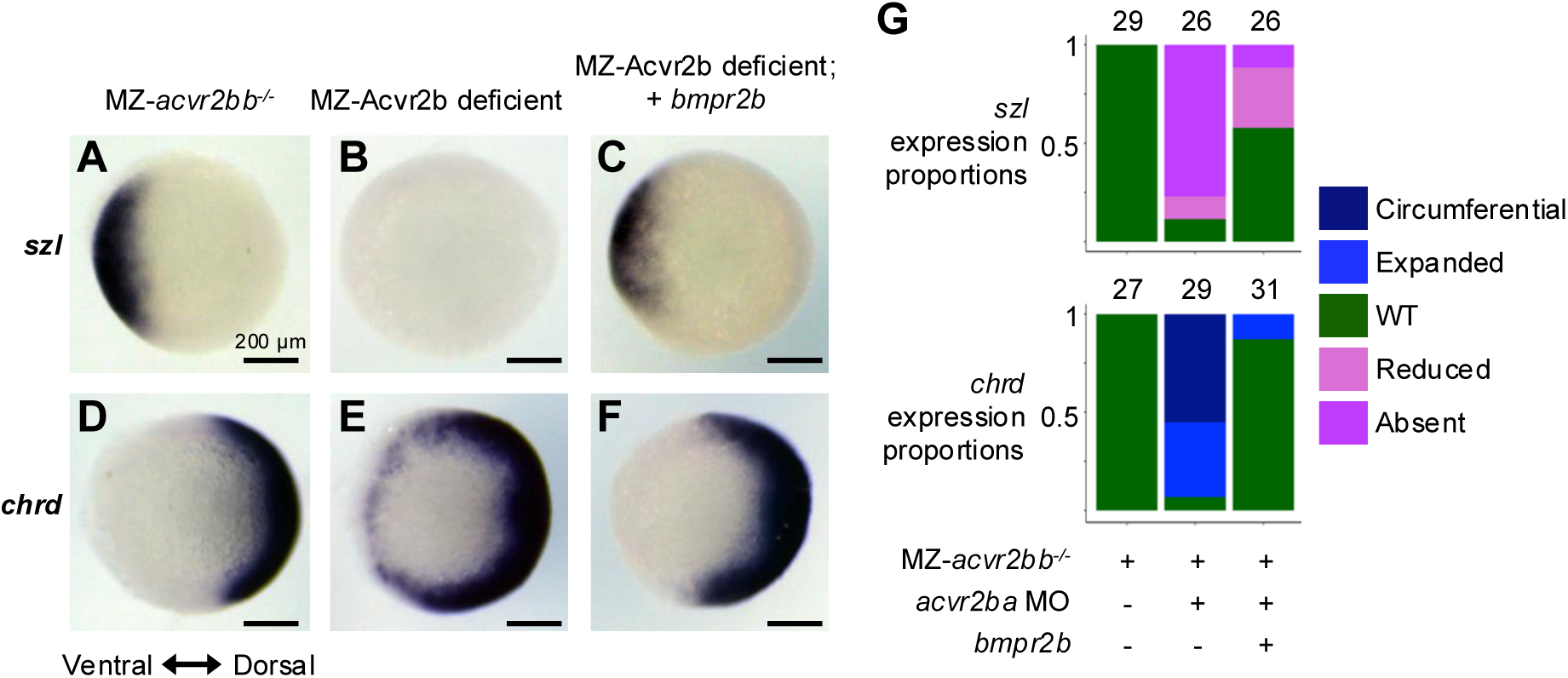
Bmpr2 receptors can function redundantly to Acvr2b in DV patterning. Representative *in situ* hybridization for *szl* (A-C) and *chrd* (D-F) in early gastrula (60% epiboly) stage embryos. Ventral *szl* expression (A) is absent in MZ-Acvr2b deficient embryos (B) and rescued by expression of *bmpr2b-venus* (C). Similarly, dorsal *chrd* expression (D) is circumferentially expanded in MZ-Acvr2b deficient embryos (E) and rescued by expression of *bmpr2b-venus* (F). Proportion of embryos with observed expression patterns of *chrd* (top) and *szl* (bottom) (G). Number of embryos assayed above bar from 4 experiments. Scale bars are 200 μm.

### ACVR1-R206H requires maternal and zygotic contributions of both Acvr2ba and Acvr2bb to signal

A recent study found that MO knockdown of both Acvr2ba and Acvr2bb in zebrafish reduced signaling by the Fibrodysplasia Ossificans Progressiva (FOP) disease receptor, ACVR1-R206H, that causes ectopic bone formation in humans (Lalonde et al., 2023); however, knockdown of Acvr2b did not completely abrogate ectopic signaling as expected if these receptors mediate all BMP signaling in this context. Given our finding that maternal and zygotic Acvr2b receptor deficiency abrogates BMP signaling, we asked if they were similarly required for ectopic ACVR1-R206H activity. We expressed *ACVR1-R206H* mRNA in either wild-type embryos (Figure 8A,B,G columns 1-2), combinations of MZ-*acvr2* mutant alleles (Figure 8C-F,G columns 7-12), or exclusively in zygotic *acvr2* mutant embryos (Figure 8G columns 3-6). We found that ACVR1-R206H ventralized wild-type embryos, as expected, as well as zygotic *acvr2* mutant embryos (Figure 8A,B,G columns 1-6). Interestingly, MZ loss of either Acvr2ba or Acvr2bb abrogated ventralization (Figure 8C-F,G columns 7-12), indicating that reduced Acvr2b receptor pools restrict *ACVR1-R206H* signaling. In support of this, *ACVR1-R206H* expression in MZ-*acvr2bb^−/−^*embryos with completely wild-type maternal and zygotic contributions of Acvr2ba also prevented ventralization/ectopic signaling, indicating that loss of just one Acvr2b receptor limits ACVR1-R206H hyperactivity (Figure 8G, columns 11-12). MZ-*acvr2bb^−/−^* embryos (Figure 8G column 11) are phenotypically wild-type, indicating that Acvr2ba, which remains present, is sufficient for normal BMP signaling but restrictive for ACVR1-R206H ectopic signaling. These results support a model in which Acvr2b receptors are limiting for FOP ACVR1-R206H hyperactivity in a dose-dependent manner.

**Figure 8:**
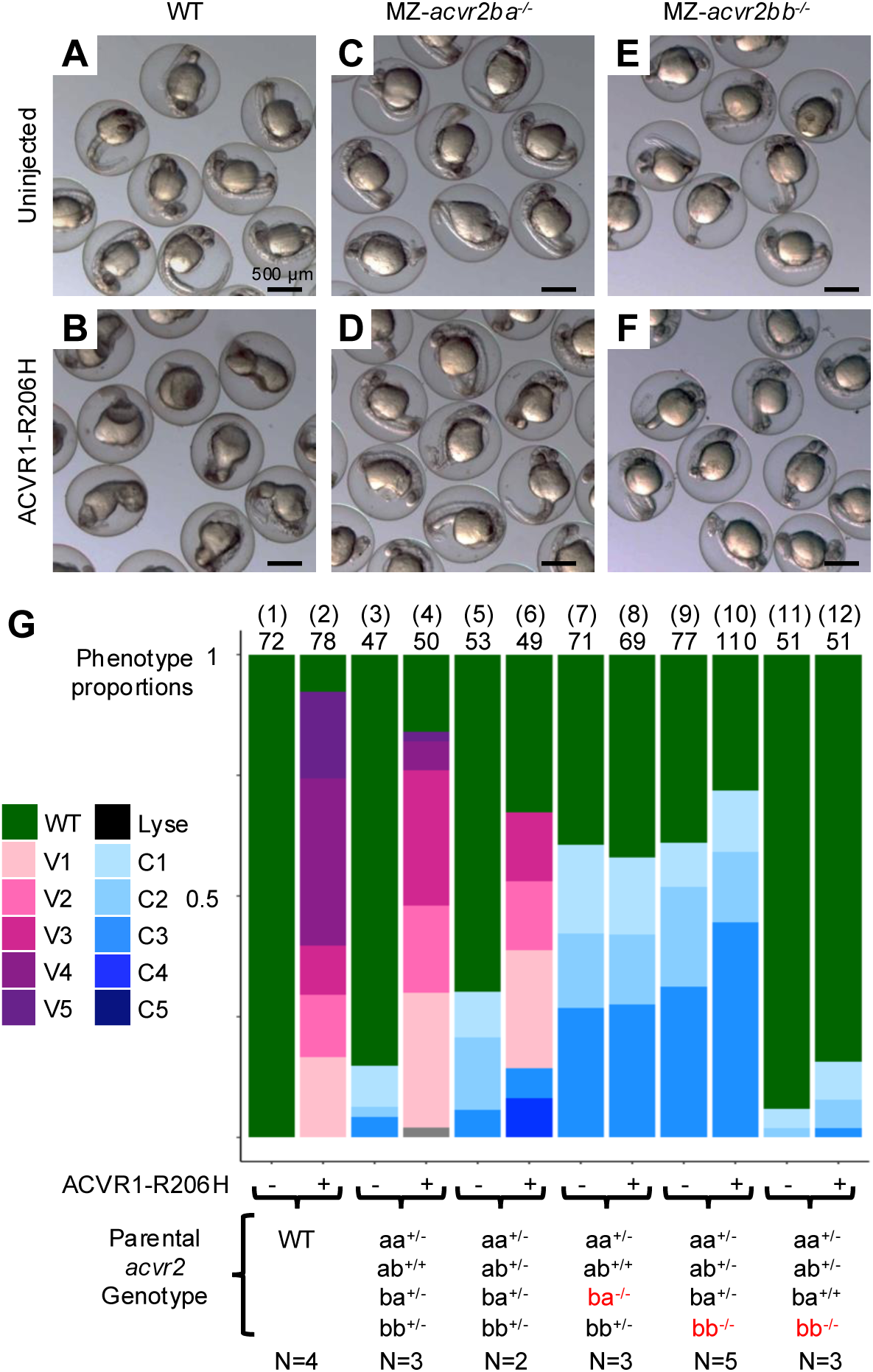
Dose dependent requirement for Acvr2b receptors in FOP ACVR1 signaling. Representative images of 1 dpf embryos from wild type (WT) (A) and MZ-*acvr2ba* and MZ-*acvr2bb* embryos displaying mild dorsalization (C,E). Injection of ACVR1-R206H severely ventralizes WT embryos (B) but does not change the phenotypes of MZ-*acvr2ba* or MZ-*acvr2bb* embryos, indicating a failure to induce hyperactive signaling without the maternal and zygotic pools of either receptor. (G) Proportions of embryos displaying different phenotypes with or without ACVR1-R206H expression from different crosses, with maternal-zygotic mutant alleles highlighted in red. N indicates total number of individual crosses examined, number above bar indicates number of individual embryos examined, number in parentheses identifies column. Scale bars are 500 μm.

## Discussion

Here we show that both maternal and zygotic contributions of Acvr2ba and Acvr2bb play major roles in transducing BMP signaling in DV axial patterning during gastrulation (Figure 2,3). MZ-Acvr2b deficient embryos display a loss of pSmad5 at early gastrula stages and circumferential expansion of dorsal fates at the expense of ventral cell fates (Figure 4, 5). Our data indicate that Acvr2b receptors are the predominate type II receptors mediating BMP signaling during DV patterning in the gastrula, and that endogenous Acvr2a levels are insufficient for patterning, but may be sufficient to drive limited signaling in the ventral-animal pole (Figure 4H, 5). Thus, the BMP signaling mechanism in the zebrafish gastrula is comprised of a Bmp2/7 heterodimer that signals through a receptor tetramer consisting of Acvr1l, Bmpr1a, and a pair of Acvr2b receptors, with Acvr2a receptors playing a minor role in signaling.

Type II Acvr2 receptors are shared by both BMP and Nodal signaling, transducing signaling through Smad5 and Smad2/3, respectively (reviewed in Zinski et al., 2018). Surprisingly, zygotic F0 CRISPants of all four Acvr2 genes show no reduction in the BMP driven pSmad5 gradient in the early gastrula, while Nodal induced pSmad2/3 is reported to modestly increase in an unknown manner (Preiß et al., 2022). We find that maternal and zygotic loss of Acvr2b genes leads to complete embryonic dorsalization (Figure 4), in line with loss of BMP signaling, but observe no consistent morphological phenotypes indicative of Nodal loss. The absence of Nodal related phenotypes in MZ-*acvr2aa^−/−^*;*acvr2ab^−/−^*embryos shows that Acvr2b receptors can transduce Nodal signaling in addition to BMP signaling, supporting a model in which both Acvr2a and Acvr2b receptors signal redundantly in these pathways in the zebrafish. Hence, maternal and zygotic depletion of all Acvr2 genes may be needed to eliminate early Nodal signaling in the embryo.

Interestingly, our data support that limited Acvr2a receptors in MZ-Acvr2b deficient embryos function in Nodal signaling at the expense of BMP signaling (Figure 6). We found that loss of Nodal signaling in MZ-Acvr2b deficient embryos was sufficient to allow BMP signaling at the embryonic margin, suggesting that competition for receptor binding can drive differential signaling outcomes *in vivo*, providing a possible model for expanded Nodal signaling in zygotic Acvr2 CRISPants (Preiß et al., 2022). Why Acvr2a preferentially signals through Nodal over BMP is unclear. One possibility is that Nodal binds Acvr2a with higher affinity than Bmp2 or Bmp7. Surface plasmon resonance (SPR) studies suggest that Nodal displays weak affinity for ACVR2A or ACVR2B, while BMP7 displays higher affinity for both receptors (Aykul et al., 2015; Aykul and Martinez-Hackert, 2016; Chu et al., 2022). However, *in vivo* fluorescence correlation microscopy in zebrafish shows that the Nodal proteins, Squint and Cyclops, bind to Acvr2b with nanomolar affinity (Wang et al., 2016). This discrepancy likely results from SPR experiments lacking the Nodal TDGF1 (Oep) co-receptor, which is required to assemble the Nodal signaling complex (Yeo and Whitman, 2001) and thus is essential to Nodal signaling in early development (Ding et al., 1998; Gritsman et al., 1999). As such, further studies comparing the *in vivo* receptor binding affinities for Nodal and BMP ligands are needed to determine if ligand-receptor preference is the source of signaling bias in this context.

Recent studies have shown that BMP and Nodal type I receptors compete for type II receptor binding in preformed complexes, which could drive the margin specific loss of pSmad5 in MZ-Acvr2b deficient embryos. Preformed receptor complexes are non-signaling complexes in which type I and II receptors form ligand-independent dimers (Agnew et al., 2021; Gilboa et al., 2000; Nohe et al., 2002). In cell culture, the BMP type I receptor ACVR1 and the Activin/Nodal-specific type I receptor ACVR1B compete to associate with ACVR2A in a preformed complex, with ACVR1B displaying a stronger affinity for ACVR2A than ACVR1 (Szilágyi et al., 2022). Whether ACVR1 or ACVR1B complex with ACVR2A determines if SMAD1/5/9 or SMAD2/3 is activated, respectively (Szilágyi et al., 2022). In the zebrafish, the Nodal type I receptor, Acvr1ba, is restricted to the embryonic margin (Preiß et al., 2022; Renucci et al., 1996). As such, it is possible that when Acvr2 receptors are sparse, as in MZ-Acvr2b deficient embryos, Nodal type I receptors preferentially complex with Acvr2a at the margin and restrict BMP signaling in these cells.

We previously demonstrated that Acvr1l kinase activity is necessary for signaling in the gastrula, though the kinase activity of the other required type I receptor, Bmpr1a, is dispensable (Tajer et al., 2021). Why only one receptor kinase acts in signaling when two receptors are available, and what facilitates the sub-functionalized kinase activity through Acvr1l in this context, remain important questions to understand BMP signaling. One possibility is that Acvr2b receptors can only phosphorylate and activate Acvr1l in the signaling complex, leaving Bmpr1a unphosphorylated. In this case, Bmpr1a may require the alternative type II receptor class, Bmpr2, for activation. If so, introducing Bmpr2 could be sufficient to activate the Bmpr1a kinase. Our results show that Bmpr2b can signal in the absence of Acvr2b (Figure 7), though our experiments do not distinguish if this receptor complex requires both Acvr1l and Bmpr1a, as in endogenous signaling. BMPR2 exhibits unique type I receptor interactions in different contexts, so ectopic Bmpr2b in the gastrula could signal via different models. For example, in myeloma cells, BMPR2 antagonizes ACVR1 phosphorylation of SMAD1/5/9 phosphorylation, which is proposed to result from BMPR2 inhibiting ACVR2-ACVR1 complex formation (Olsen et al., 2018). In contrast, BMPR2 in gonadotrophic cells promotes signaling through either ACVR1 or BMPR1A, though if these receptors form a type I heteromeric complex in this process is not known (Ho and Bernard, 2009; Lee et al., 2007). As such, future work will need to not only determine if Bmpr2b signals through a heteromeric type I complex, but also determine the factors permitting specific type I-type II receptor combinations to signal in different contexts.

Hyperactive signaling via the human FOP disease-causing ACVR1-R206H further underscores the importance of Acvr2b receptors for BMP signaling. Maternal-zygotic loss of either Acvr2ba or Acvr2bb rendered ACVR1-R206H unable to signal ectopically, even when all three other Acvr2 receptors were present. This contrasts with Acvr2ba and Acvr2bb in BMP signaling, where loss of both receptors is necessary to abrogate signaling during DV patterning. These findings are consistent with a dosage model in which reduced Acvr2b restricts ectopic signaling by ACVR1-R206H but is sufficient for wild-type Acvr1l signaling. Further, a recent study found that preformed ACVR2B homodimers increase the ability of ACVR1-R206H to form ligand-independent signaling oligomers (Szilágyi et al., 2024), consistent with our finding that ACVR1-R206H hyperactivity is restricted when the amount of Acvr2b receptors is reduced.

In all, our findings reveal a key function for Acvr2ba and Acvr2bb in mediating BMP signaling during DV axial patterning in the zebrafish gastrula. Importantly, we find dose-dependent requirements for Acvr2 in activating BMP signaling specifically at the embryonic margin and for ectopic disease signaling by ACVR1-R206H. It will be important to next determine if limited type II receptors drive competition between Nodal and BMP signaling in embryonic signaling and further interrogate how type II receptors facilitate Nodal mesendoderm induction and patterning. Additional studies will need to determine if and how Acvr2b regulates sub-functionalized type I receptor kinase activity, either by selective phosphorylation of Acvr1l or through interactions with other factors that delineate exclusive kinase activity through Acvr1l. Further exploration of how type II receptors mediate BMP signaling in other signaling contexts are needed to fully understand how the receptor landscape drives interpretation of key signaling pathways for embryonic patterning.

## Author Contributions

**Conceptualization**: J.H.P., B.T., M.C.M.; **data curation**: J.H.P., B.T., J.W.; **formal analysis**: J.H.P., B.T., M.C.M.; **funding acquisition**: J.H.P., M.C.M.; **investigation**: J.H.P., B.T., J.W., F.W.; **methodology**: J.H.P., B.T., J.W.; **project administration**: M.C.M.; **resources**: M.C.M.; **supervision**: M.C.M.; **validation**: J.H.P., M.C.M.; **visualization:** J.H.P., B.T.; **writing, original draft**: J.H.P., M.C.M; **writing, review & editing**: J.H.P., B.T., F.W., M.C.M.

## Declaration of Interests

The authors declare no competing interests.

## Acknowledgements

This study was supported by NIH Grant R35-GM131908 to M.C.M, NIH Grant F32-HD114468 to J.H.P, and J.W. was supported by NIH Grant R25-GM071745. We thank the University of Pennsylvania Department of Cell and Developmental Biology Microscopy Core for guidance and use of the Ziess LSM880 confocal and Ari Geller for comments on this manuscript.

## Methods

### Zebrafish husbandry

The research performed here was approved by the University of Pennsylvania Animal Care and Use Committee. Adult fish were housed at 28°C in a facility on a 13/11 light/dark cycle and cared for in accordance with institutional regulatory standards. Embryos were collected from crosses within 15 minutes of fertilization and reared in E3 media (4 mM NaCl, 0.17 mM KCl, 0.33 mM CaCl2, and 0.33 mM MgSO4) at 28-31.5°C. DNA from adult fin clips or embryos was extracted via HotShot and used for genotyping (Meeker et al., 2007).

### Phenotypic evaluation

Embryonic phenotypes were assessed at 1 dpf based on previously established criteria (Kishimoto et al., 1997; Mullins et al., 1996; Nguyen et al., 1998). Embryos were imaged in E3 using a Leica IC80HD camera and images were processed in FIJI (Schindelin et al., 2012). In brief, C1 embryos partially lack the ventral tail fin, C2 do not have a ventral tail fin nor ventral tail vein, C3 embryos lack a yolk extension, C4 embryos have fewer than 13 somites evident at 1 dpf, and C5 embryos exhibit strong radial dorsal convergence, evidenced by elongation at 12 hours post fertilization and lysis by 1 dpf. For ventralized staging, V1 embryos are characterized by reduced head size, V2 lack notochords and have further reduction in head size, V3 lack anterior head structures and have slightly wider posterior somites, V4 display an enlarged yolk extension and wider somites, and V5 embryos display greatly expanded ventral mesoderm and blood islands. For genotyping, embryos were sorted by phenotype and then isolated into 96 well plates in methanol, preventing bias in scoring, before HotShot lysis to extract DNA for genotyping.

### Microinjection of mRNA and MO

Embryos were collected within 15 minutes following fertilization in E3 media and kept at 22°C during injections. For each experimental replicate, a single needle containing either mRNA or MO was prepared and each injected into embryos on the same day. For conditions with multiple injections, all needles were prepared at the same time and serially injected into embryos, allowing independent assessment of the effect of each reagent on its own. Injection solutions were prepared such that 1-2nL yielded concentrations reported in Table 1, with a final concentration of 0.1M KCl and 0.1% phenol red. mRNA was synthesized from sequenced plasmids using the Sp6 mMessage mMachine kit from Ambion. Previously published *acvr2bb* MO (Dogra et al., 2017) led to off target head necrosis at concentrations that had no other phenotypes, so we designed a new 5’UTR targeting morpholino and injected a mix of both *acvr2bb* AUG MO and 5’UTR MO to increase specificity while reducing toxicity.

**Table 1:**
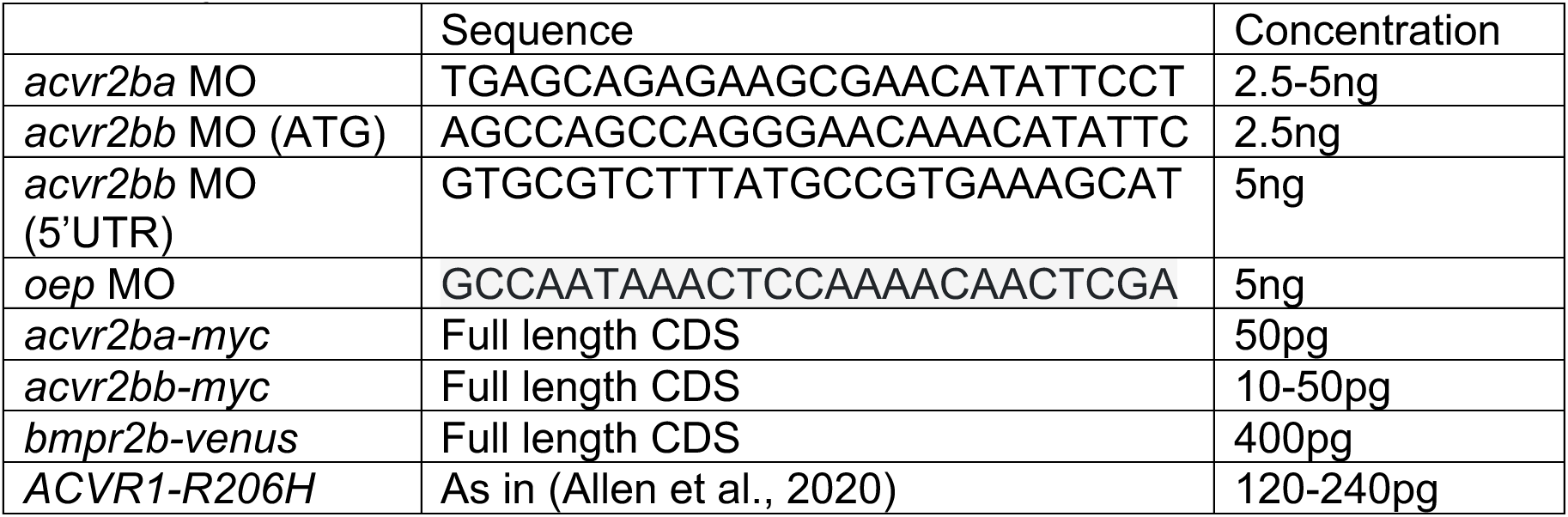
Injection constructs and concentrations.

### CRISPR/Cas9 sgRNA generation

We mutagenized *acvr2aa*, *acvr2ba*, and *acvr2bb* using CRISPR-Cas9. Targets were selected with the assistance of the now discontinued CRISPR targeting resource on (http://crispr.mit.edu/) or the web tool CHOPCHOP ((https://chopchop.cbu.uib.no) (Montague et al., 2014). *acvr2aa* and *acvr2ba* Single Guide RNAs (sgRNAs) were synthesized in-vitro from PCR amplified templates using the cloning free method used in Gagnon et al 2014 (Gagnon et al., 2014), with the variation that sgRNA templates were amplified using Phusion Polymerase (ThermoFisher F553S), using the kit protocol scaled to a reaction size of 40μL. sgRNA templates were purified using the MinElute purification kit (QIAGEN 2800) to a final volume of 20μL. sgRNAs were synthesized from templates in-vitro using the MEGAshortscript T7 kit (ThermoFischer AM1354), using 8μLs of undiluted template in a reaction volume of 20μL. MEGAshortscript reactions were run overnight as this was found to increase yield. sgRNAs were purified using ethanol precipitation, as in Gagnon et al., 2014. Purified sgRNAs were assessed and quantified by running dilution series on a glyoxal/sodium phosphate buffer RNA gel and comparing to RiboRuler (ThermoFischer SM1821). *acvr2bb* sgRNAs were ordered directly from IDT rather than synthesized from DNA oligos. sgRNA sequences are presented in Table 2.

**Table 2:**
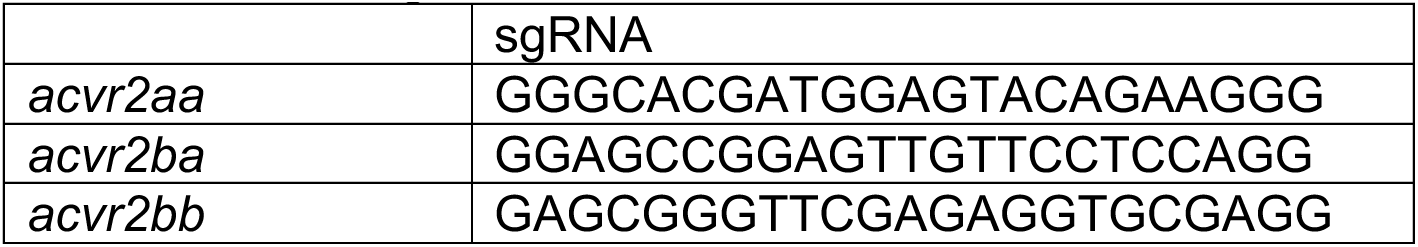
CRISPR guides.

### CRISPR injection and allele identification

Immediately prior to injection, 2.5μL of undiluted sgRNA was mixed with 1μL of 5mg/mL Cas9-NLS protein (PNA Bio CP01-50), 1μL of Phenol Red, and 0.5μL 1M KCl (total volume 5μL) and kept at room temperature. 2nL of this injection mix was injected into wild-type embryos. For *acvr2bb*, guides were injected into embryos of triple *acvr2aa*;*acvr2ab*;*acvr2ba* heterozygous incrosses. To identify mutant alleles, F1 DNA samples were cleaned with ExoSapit (AppliedBiosystems) and sequences were extracted from heterozygous trace data with the help of the web app PolyPeakParser (Hill et al., 2014).

### Genotyping

Kompetitive allele specific PCR amplification (KASPar) genotyping (Smith and Maughan, 2015)was used to genotype *acvr2aa*, *acvr2ab*, and *acvr2bb* (Table 3). A PCR band size difference was used to genotype *acvr2ba* mutants (162 bp wild-type band and 241 bp mutant band) using the standard NEB 2xTaq protocol modified to include 3% DMSO (12.5μL 2xTaq, 1 μL 10μΜ forward and reverse primer, 0.75μL DMSO, 5.25μL water, 5μL gDNA). Primers used for genotyping *acvr2aa*, *acvr2ba*, and *acvr2bb* are reported in Τable 3.

**Table 3:**
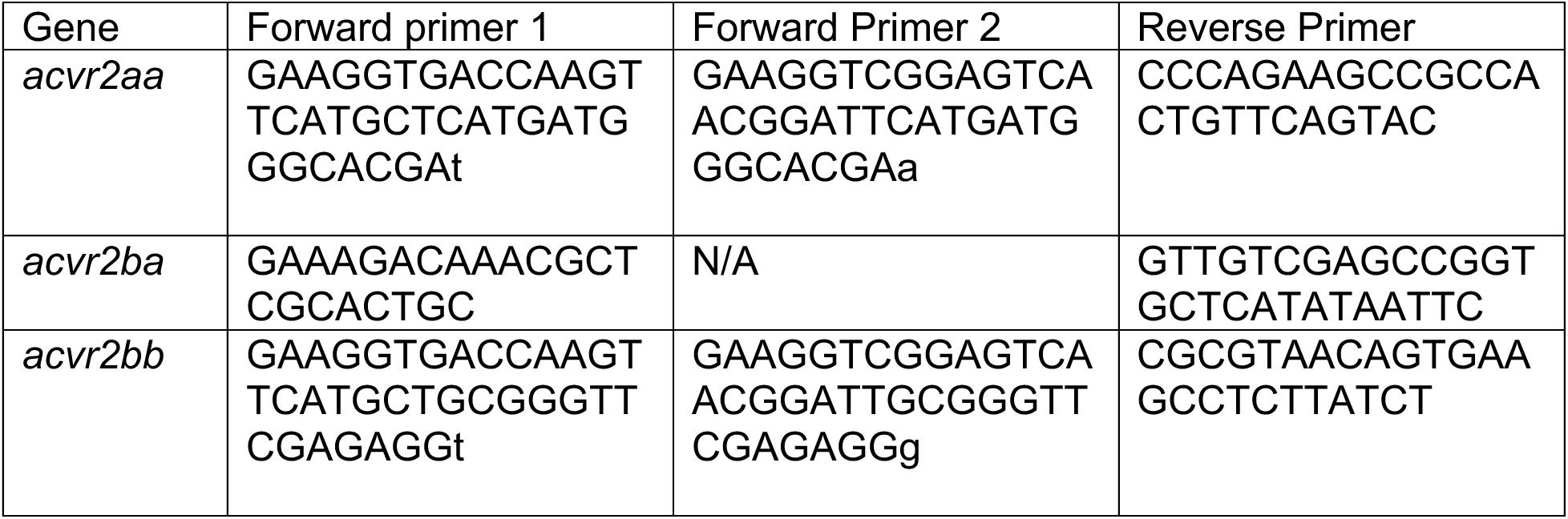
Genotyping primers.

For *acvr2ab*, the sequence provided to LGC genomics was: TGGAGAAAAGGACAAGCGRCGACACTGTTTCTCCACCTGGAAGAACCGCT[C/A]GG GRACCATCGAGATGGTSAAGCAGGGCTGCTGGCYGGACGATGTCAACT.

### Immunofluorescent staining and quantification

pSmad5 immunostaining and quantification was performed as described (Zinski et al., 2019). Briefly, shield stage embryos were fixed in 4% formaldehyde in PBS O/N at 4°C, washed 3×20 minutes in PBS with 0.1% TritonX (PBST) and deyolked manually, then incubated in blocking solution (10% FBS, 1% DMSO in PBST). Samples were then incubated O/N at 4°C with a 1:200 dilution of rabbit pSmad1/5/9 antibody (Cell Signaling 13820) in block, washed 3×20 minutes with PBST, then incubated O/N at 4°C in 1:500 dilution of anti-rabbit Alexa 647 (Invitrogen-21245) and 1:2000 SYTOX green (Fisher S7020) in block, followed by 3 to 4, 20 minutes PBST washes and 1 O/N wash at 4°C. Immunostained embryos were then dehydrated in methanol and cleared using a 1:2 ratio of benzyl alcohol (Sigma B-104) and benzyl benzoate (Sigma B-6630) to match the refractive index of yolk. Embryos were mounted animal pole down, with the DV axis parallel to the cover slip and imaged using a Ziess LSM880 confocal with an LD LCI Plan-Achromat 25x/0.8 lmm Corr DIC M27 multi-immersion objective set to oil-immersion.

Image files were converted to TIFFs and nuclei were segmented by Sytox staining using a custom MATLAB script (Zinski et al., 2017). Then, pSmad5 intensity for each nucleus was calculated and resulting embryos were aligned across the DV axis and aligned to a reference WT embryo. To normalize staining and imaging differences across experiments, the mean of each WT embryo was calculated, and an average of all WTs per experiment was determined. Then, these means were used to determine a ratio relative to a reference experiment to get a scaling value between experiments, which was applied to all embryos stained and imaged from that single experiment. Population means were generated across each experimental group and nuclei were projected into equilateral triangles, tiling the embryo to generate heatmaps. Margin gradients were calculated by examining a 20μm band of cells at the indicated position in 5 degree increments (Zinski et al., 2019). To assess significance, a two-tailed t-test was performed for each bin and adjusted p-values (q-values) were determined using a Benjamini-Hochberg correction. For ventral cell comparison in Figure 6, cells in a 20° subset of the band were averaged per embryo, then a single mean per replicate was generated. A paired, two-tailed t-test was performed between each condition and p-values were adjusted (q-value) using the Holm-Bonferroni method. Representative images were processed using FIJI, taken from the same experiment and processed using identical parameters in each figure.

### in situ hybridization

Whole mount *in situ* hybridization was performed on shield, 60%, 1-3 somite, and 10-somite stage embryos using DIG labeled anti-sense probes (made with the Roche 11277073910 kit), following the multi-basket protocol established in (Thisse and Thisse, 2008) with some modifications. Briefly, no proteinase K was added at any stage, probes were incubated overnight at 65°C. Blocking and antibody incubation were performed with blocking reagent (Roche, 11096176001) diluted in maelic acid buffer (MAB, 100 mM maelic acid, 150 mM NaCl, adjusted to pH 7.5) after which samples were given 3 quick washes and 3, 20 minutes washes in MAB-Tw (MAB+0.1% Tween-20). Samples were then washed 2 times for 10 minutes in PBS with 0.1% Tween-20 (PBS-Tw) before developing in BM-Purple (Roche, 11442074001). The reaction was quenched with 3-5 washes in PBS-Tw. Probes for *pax2.1* (Krauss et al., 199*2)*, *krox20* (Oxtoby and Jowett, 1993), *szl* (Yabe et al., 2003), and *chrd* (Miller-Bertoglio et al., 1997) were generated previously, while *bmpr2b* was made from a full length CDS plasmid. Probes for *bmpr2a* were synthesized from genomic DNA PCR products using a primer set containing a T7 sequence: F – CTCAGAGTCCTGCAATGCCA, R – taatacgactcactatagggGTCAAGTCCTCGTTGGCTGA.

### Fluorescent in situ hybridization and quantification

Fluorescent *in situ* hybridization was performed following the same *in situ* protocol as above until antibody incubation. Samples were incubated in anti-DIG-Horseradish Peroxidase (Roche, 11207733910, 1:1000 dilution) in blocking reagent (Roche, 11096176001), washed in MAB-Tw then PBS-Tw, and then developed for 1 hour in a 1:50 dilution of TSA Plus Cyanine 3 (Quanterix, NEL744001KT) before washing in PBS-Tw. Nuclei were stained with a 1:2000 dilution of Sytox green. Clearing, mounting, and imaging were performed as described in the immunostaining method. Analysis was performed as with pSmad5, except fluorescent intensity was extracted from a sphere of approximately 10 μm in diameter (as opposed to ∼4.5 μm for nuclear pSmad5) to account for signal in the cytoplasm. Probes for *foxi1* (Kwon et al., 2010) and *bambia* (Greenfeld et al., 2021) were generated previously.

### Analysis of published datasets

RNA-seq data from (White et al., 2017) was downloaded from https://www.ebi.ac.uk/gxa/experiments/E-ERAD-475/Results by searching for Gene ID and exporting transcripts per million (TPM) at each developmental stage. The heatmap in Figure 1 was generated from TPM data using the R package, pheatmap (Raivo Kolde, 2010).

### Generation of plots

Stacked boxplots were generated using ggplot2 (Wickham, 2016). pSmad5 intensity and margin gradient plots were generated using a published MatLab analysis pipeline (Zinski et al., 2019).

## Notes

### Competing Interest Statement

The authors have declared no competing interest.

### Summary of Updates

This revision includes additional data figures and research data not present in the original publication, with major addition in figure 5 and new figures 6 and 7.

