## Supplemental Figures for "Acvr2b receptors dose-dependently regulate dorsoventral patterning, FOP ACVR1-R206H signaling, and limit BMP/Nodal signaling"

### Figure S1: Acvr2 mutants result in predicted frameshifts with premature stop codons

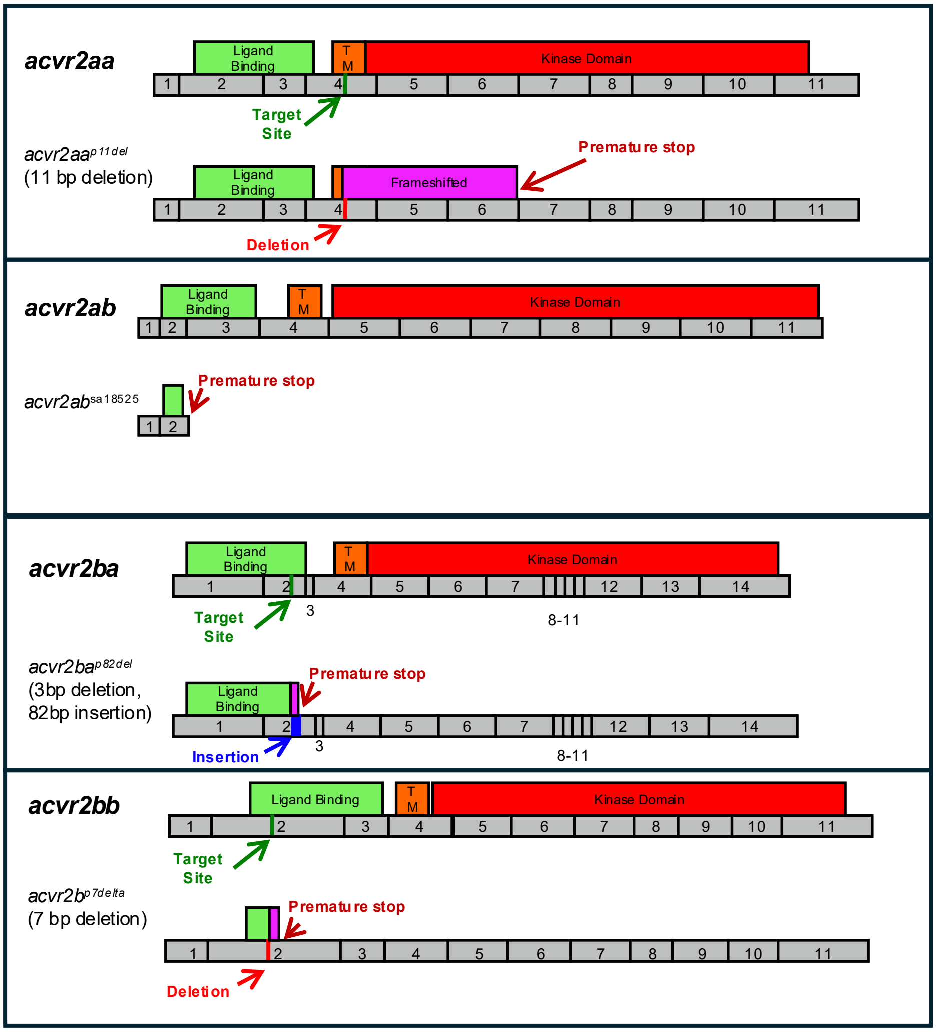


**Figure S1: Acvr2 mutants result in predicted frameshifts with premature stop codons**
List of *acvr2* alleles generated with CRISPR or obtained from ZIRC, and their positions relative to the CRISPR target site (green), exon structure (grey), and major protein domains: Ligand binding domain (light green), transmembrane domain (orange), and kinase domain (red). Insertions shown in blue. Presumptive modifications to the transcript or protein are represented in purple.

#### Figure S2: Maternal loss of Acvr2ba and Acvr2bb enhances patterning defects

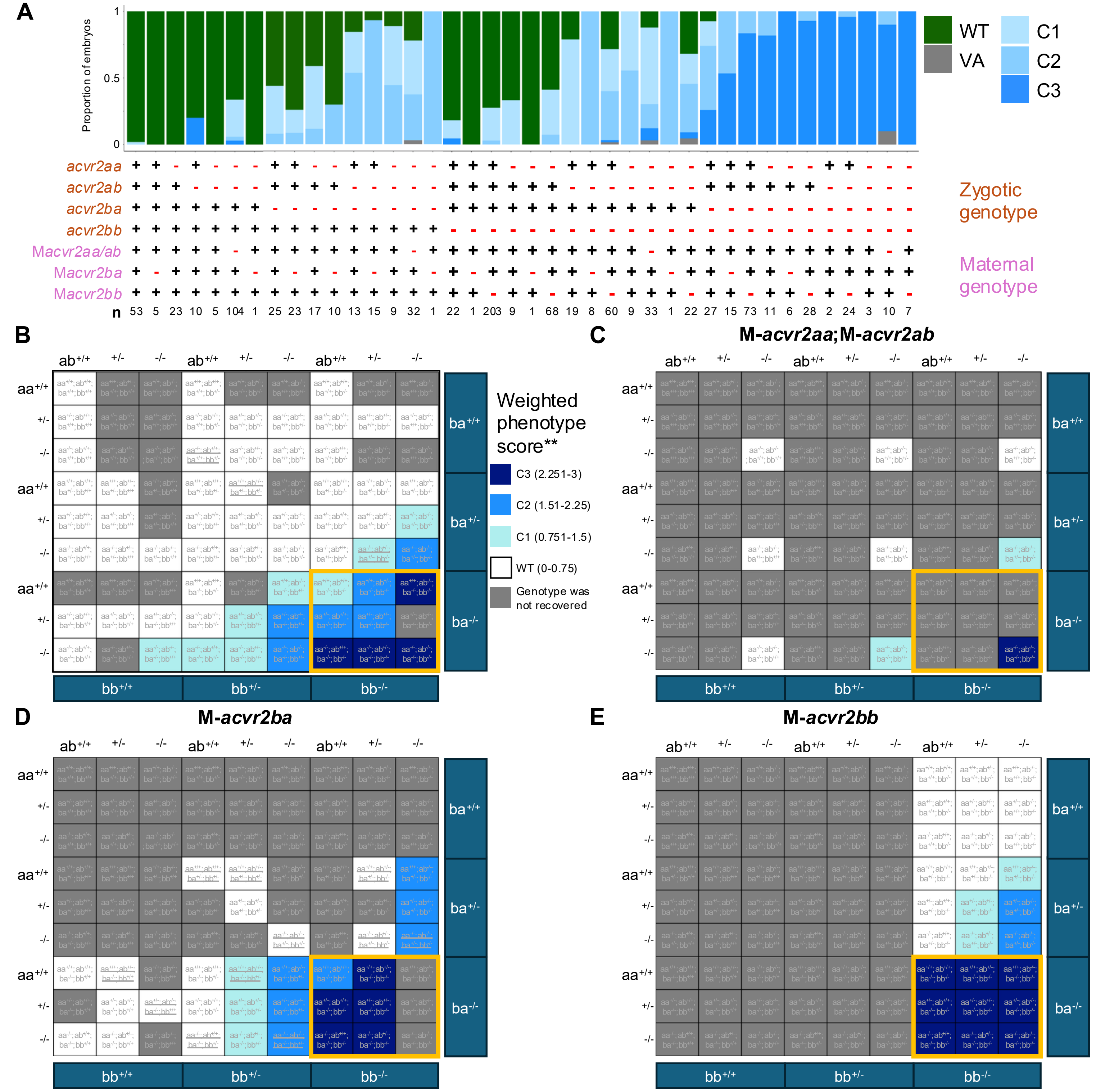


**Figure S2: Maternal loss of Acvr2ba and Acvr2bb enhances patterning defects**

A) Representative phenotypes distribution for combinations of maternal and zygotic *acvr2* mutants from Figure 2. As these crosses generate 81 distinct zygotic genotypes and 4 different maternal genotypes, we narrow our analysis by combining zygotic WT and heterozygous genotypes (+) vs mutant (-). Not all phenotypically wild-type embryos were genotyped, so these proportions represent possible phenotypic outcomes. n indicates number of individual embryos represented. B-E) Weighted dorsalization scores for each genotype based on assayed embryos. **Not every phenotypically wild-type embryo was assayed, so in some cases these scores are skewed by low n. Phenotypically WT embryos are scored 0, with dorsalized C1, C2, and C3 embryos scored as 1, 2, or 3, respectively. Colors are binned by average phenotype score and grey indicates no embryos of such genotype(s). Maternal genotypes were: no homozygous maternal mutations (B), M-*acvr2aa*$^{-/-}$;M-*acvr2ab*$^{-/-}$ (C), M-*acvr2ba*$^{-/-}$ (D) M-*acvr2bb*$^{-/-}$ (E). Orange boxes highlight *acvr2ba*$^{-/-}$;*acvr2bb*$^{-/-}$ double mutants. Underlined genotypes indicate only 1 individual was recovered for a genotype. Parental genotypes for crosses are as follows (female X male genotype):

B) *acvr2aa*$^{+/-}$;*ab*$^{+/+}$;*ba*$^{+/-}$;*bb*$^{+/-}$ X *acvr2aa*$^{+/-}$;*ab*$^{+/+}$;*ba*$^{+/-}$;*bb*$^{+/-}$ ;
*acvr2aa*$^{+/-}$;*ab*$^{+/-}$;*ba*$^{+/-}$;*bb*$^{+/-}$ X *acvr2aa*$^{+/-}$;*ab*$^{+/-}$;*ba*$^{+/-}$;*bb*$^{+/-}$ ;
*acvr2aa*$^{+/-}$;*ab*$^{+/-}$;*ba*$^{+/-}$;*bb*$^{+/-}$ X *acvr2aa*$^{+/-}$;*ab*$^{+/-}$;*ba*$^{+/-}$;*bb*$^{-/-}$ OR *acvr2aa*$^{+/-}$;*ab*$^{+/-}$;*ba*$^{+/-}$;*bb*$^{+/-}$ X *acvr2aa*$^{-/-}$;*ab*$^{-/-}$;*ba*$^{+/-}$;*bb*$^{+/-}$ ;

C) *acvr2aa*$^{-/-}$;*ab*$^{-/-}$;*ba*$^{+/-}$;*bb*$^{+/-}$ X *acvr2aa*$^{-/-}$;*ab*$^{-/-}$;*ba*$^{+/-}$;*bb*$^{+/-}$ ;

D) *acvr2aa*$^{+/-}$;*ab*$^{+/-}$;*ba*$^{-/-}$;*bb*$^{+/-}$ X *acvr2aa*$^{+/-}$;*ab*$^{+/-}$;*ba*$^{-/-}$;*bb*$^{+/-}$ OR *acvr2aa*$^{+/-}$;*ab*$^{-/-}$;*ba*$^{+/-}$;*bb*$^{-/-}$ ;
*acvr2aa*$^{+/-}$;*ab*$^{+/+}$;*ba*$^{-/-}$;*bb*$^{+/-}$ X *acvr2aa*$^{+/-}$;*ab*$^{+/-}$;*ba*$^{+/-}$;*bb*$^{-/-}$ ;
*acvr2aa*$^{+/-}$;*ab*$^{+/+}$;*ba*$^{-/-}$;*bb*$^{+/+}$ X *acvr2aa*$^{+/-}$;*ab*$^{+/-}$;*ba*$^{-/-}$;*bb*$^{+/+}$ ;

E) *acvr2aa*$^{+/-}$;*ab*$^{+/-}$;*ba*$^{+/-}$;*bb*$^{-/-}$ X *acvr2aa*$^{+/-}$;*ab*$^{+/-}$;*ba*$^{+/-}$;*bb*$^{-/-}$ ;
